# Dissecting the genetic basis of drought escape across multiple traits in colonizing *Arabidopsis thaliana* lineages

**DOI:** 10.1101/2025.04.07.647068

**Authors:** Ahmed F. Elfarargi, Elodie Gilbault, Nina Döring, Herculano Dinis, Andreas P. M. Weber, Olivier Loudet, Angela M. Hancock

## Abstract

Drought response in plants is complex, involving integration across a range of physiological processes. However, our knowledge of how different aspects of drought response are linked at the genetic level is limited. We investigated multi-trait adaptation in *Arabidopsis thaliana* from the Cape Verde Islands (CVI). Using a high-throughput phenotyping platform that minimizes spatial heterogeneity, we measured variation in rosette area, growth rate, leaf color, water-use efficiency (WUE), and stomatal patterning under precisely controlled water conditions. Relative to the Moroccan outgroup, CVI populations evolved to be smaller rosette size with faster growth and reduced WUE, consistent with drought escape adaptation.

Genome-wide association mapping revealed evidence for pleiotropy involving *MPK12* (WUE, rosette area, growth rate, and leaf color), *NHL26* (WUE and leaf color), *SUVH4* (stomatal patterning, rosette area, and leaf color), and *FRI* (WUE and leaf color), along with an enrichment of signals in ABA response. This study advances our knowledge of the genetic mechanisms driving plant adaptation to novel precipitation environments. By identifying key genetic components and their contributions to multi-trait adaptation, our findings offer insights into how plants respond to environmental challenges and contribute to predicting plant responses to future climate change.

## Introduction

Plants are directly exposed to environmental stress and must evolve mechanisms to overcome such challenges (Chaves *et al*., 2003; Lawlor, 2013). Among these stresses, water availability stands out as a critical factor that limits plant growth and productivity (Boyer, 1982) and profoundly influences development, morphology, and physiology (Stebbins, 1952; Bohnert *et al*., 1995). In diverse environments, populations exhibit a broad spectrum of phenotypic variation, which is essential for survival and adaptation. This variation often arises from phenotypic plasticity, the ability of a single genotype to exhibit different phenotypes in response to environmental changes (Sultan, 2000; Gratani, 2014; Josephs, 2018). Plasticity enables plants to rapidly adjust to fluctuating conditions, such as changes in water availability (Pigliucci, 2005). However, the underlying genetic architecture plays a pivotal role in shaping long-term evolutionary responses to stress. To optimize growth and fitness under water-limited conditions, plants have evolved three primary strategies: drought tolerance, drought avoidance, and drought escape (Ludlow, 1989). Drought tolerance maximizes water-use efficiency (WUE) by balancing carbon assimilation and water loss via transpiration (Scott, 2000). Drought avoidance enhances water uptake through the development of extensive root systems and minimizes water loss by reducing transpiration (Levitt, 1972). In contrast, drought escape relies on accelerated flowering to complete the life cycle before the onset of severe drought (McKay *et al*., 2003; Sherrard & Maherali, 2006). These strategies, involving intricate control of physiological processes, and morphological and developmental traits, reflect a dynamic interplay between phenotypic plasticity and the genetic networks that govern plant adaptation.

One critical aspect of this genetic interplay is pleiotropy, where a single gene influences multiple traits, linking diverse responses within a shared genetic framework (Paaby & Rockman, 2013). For instance, traits associated with drought escape, such as accelerated flowering, may be genetically correlated with traits like growth rate or WUE, reflecting trade-offs or coordinated adaptations to water stress (Lande, 1980; Mackay & Anholt, 2024).

Depending on the nature of these genetic linkages, pleiotropy can either facilitate adaptation through synergistic effects where traits align to meet environmental demands or constrain it when antagonistic effects cause conflicts between traits (Zhang, 2023). Understanding how pleiotropy shapes these coordinated responses is crucial for predicting plant evolutionary trajectories under changing climates.

Among the strategies plants use to cope with water stress, plants that rely on drought escape prioritize rapid growth and reproduction, emphasizing the need to balance immediate survival with overall fitness (Ludlow, 1989). The ability of plants to respond to drought conditions relies on a complex genetic network that integrates hormonal signaling pathways, including abscisic acid (ABA), ethylene, and auxin, which coordinate key physiological and developmental processes (Raghavendra *et al*., 2010; Peleg & Blumwald, 2011). ABA plays a central role in drought responses by activating cascades of drought-responsive genes that trigger morphological and physiological changes, such as stomatal closure and reduction in leaf size, resulting in an overall enhancement in WUE (Abe *et al*., 2003; Umezawa *et al*., 2010). However, these responses are shaped by contrasting selection pressures and physiological constraints, resulting in trade-offs between growth and water conservation. For instance, pleiotropic effects can link genes regulating stomatal traits and WUE to flowering time and biomass allocation, reflecting the interconnected nature of adaptive traits (Verslues *et al*., 2006; Tuteja, 2007). These responses reveal the trait complexity and the trade-offs that plants face to balance growth and survival. Understanding these complex genetic relationships and trade-offs is crucial for unraveling the genetic basis that underpins plant adaptation to drought, the sophisticated strategies plants use to cope with arid conditions, and the co-evolution of such traits in response to environmental pressures.

Natural populations of *Arabidopsis thaliana*, the major molecular model organism for higher plants, thrive in diverse habitats with considerable variation in water availability and other environmental factors (Alonso-Blanco & Koornneef, 2000; Koornneef *et al*., 2004; Mitchell-Olds & Schmitt, 2006; Verslues & Juenger, 2011; Weigel, 2012; Assmann, 2013). Early research on *A. thaliana* emphasized the significance of natural variation within the species as a valuable tool for understanding the connection between molecular perturbations and phenotypic changes (Somerville & Koornneef, 2002). Moreover, *A. thaliana* has served as a useful model for investigating the molecular genetic basis of complex traits (Mitchell-Olds & Schmitt, 2006). Local adaptation to different environmental conditions, particularly precipitation and humidity, has influenced allele frequency distributions across its European range (Hancock *et al*., 2011; Fournier-Level *et al*., 2011; Lasky *et al*., 2012; Exposito-Alonso *et al*., 2018; Ferrero-Serrano & Assmann, 2019). Studies focusing on local population sampling have proven particularly powerful for uncovering the genetic basis of adaptation to environmental variation, offering valuable insights into evolutionary processes at fine scales (Brachi *et al*., 2011; Josephs *et al*., 2017). These approaches highlight how local populations can be used to disentangle the molecular and genetic factors underlying plant adaptation to diverse and challenging environments.

A single *A. thaliana* accession, Cvi-0, was collected in the Cape Verde Islands (CVI) 42 years ago (Lobin W., 1983) and has been studied intensively due to its divergence from other well-studied accessions across various traits, such as WUE, stomatal behavior, flooding tolerance, and flowering time (Alonso-Blanco *et al*., 1998; McKay *et al*., 2003; Juenger *et al*., 2005; Christman *et al*., 2008; Xu & Zhou, 2008; Vashisht *et al*., 2011; Monda *et al*., 2011; Kenney *et al*., 2014). Mapping studies using recombinant inbred lines (RILs) derived from crosses between Cvi-0 and Ler-0 or Col-0 have identified quantitative trait loci (QTL) associated with these divergent traits, providing valuable insights into the genetic basis of adaptive variation and revealing complex interactions between genetic and environmental factors (Alonso-Blanco *et al*., 1998; Juenger *et al*., 2005; Tisné *et al*., 2013; Marchadier *et al*., 2019).

More recently, we collected new population samples of *A. thaliana* from the islands of Santo Antão and Fogo (Fulgione *et al*., 2022). As is common for endemic plant species in Cape Verde, *A. thaliana* is found on rocky outcrops where moisture is provided mainly via humid trade-winds (Brochmann *et al*., 1997; Elfarargi *et al*., 2023). Sequencing and population genetic analysis revealed that the two CVI island populations experienced strong colonization bottlenecks, which removed nearly all pre-existing genetic variation. Consequently, genetic variation segregating within each island population is highly distinct, both between islands and in comparison to mainland population (Fulgione *et al*., 2022). Notably, the vast majority of genetic variants (99.9%) observed in these populations are attributed to novel mutations that arose post-colonization, highlighting the role of recent mutations in shaping genetic diversity in these isolated populations (Fulgione *et al*., 2022).

In the ecological context of CVI, where growing seasons are short and plants rely heavily on moisture provided by humid trade-winds rather than direct rainfall (Brochmann *et al*., 1997), selection appears to have favored a drought-escape strategy. CVI populations exhibit more open stomata, enabling higher rates of photosynthesis and faster growth to facilitate drought escape (Fulgione *et al*., 2022; Elfarargi *et al*., 2023). The high relative humidity in this environment may mitigate the physiological trade-off between photosynthesis and transpiration, thereby allowing plants to optimize carbon gain without experiencing excessive water loss (Elfarargi *et al*., 2023). However, localized variation in microclimatic conditions, including temperature and humidity, may impose heterogeneous selection pressures, potentially maintaining alleles such as *MPK12* at intermediate frequencies (Elfarargi *et al*., 2023). This highlights the role of both spatial and temporal environmental heterogeneity in maintaining functional genetic variation, illustrating how natural selection acts on trait evolution to align with variable local environmental pressures (Fulgione *et al*., 2022; Elfarargi *et al*., 2023).

In this study, we measured eight phenotypic traits in response to precisely controlled drought stress in an expanded sampling of the Santo Antǎo natural population, where the Cvi-0 accession originated. Compared to our earlier work (Elfarargi *et al*., 2023), we extended this study to incorporate Fogo and Moroccan populations while also examining a broader set of phenotypic traits, enabling a broader comparison of phenotypic and genetic variation across islands and the mainland. We found that for several traits, phenotype distributions for the CVI islands are significantly shifted compared to Morocco, which represents the closest outgroup population. We further conducted genome-wide association (GWA) mapping using univariate and multivariate GWAS models to dissect the genetic basis of phenotypic adaptation underlying drought stress in the CVI *A. thaliana* populations and to compare them to the Morocco *A. thaliana* population. This analysis revealed a moderately complex genetic architecture marked by several major effect variants. Notably, some loci identified through GWAS co-localized across traits, suggesting pleiotropic effects and their involvement in multiple drought-responsive pathways. Overall, these findings enrich our understanding of the genetic complexities of plant adaptation, reflecting the critical trade-offs plants maintain for their survival in arid environments and underlying the importance of understanding genetic correlations to predict future evolutionary dynamics under changing climatic conditions.

## Results

### Population structure based on genome-wide genetic variation

Relative to previous studies we conducted on the CVI population, we expanded our set of samples to include 15 additional *A. thaliana* accessions from new locations in Santo Antão, Cape Verde, to create a panel of 189 Santo Antão accessions (Fulgione *et al*., 2022; Elfarargi *et al*., 2023). Additionally, this study includes analyses of 119 accessions from Fogo, another island in the CVI archipelago, and a representative outgroup panel of 64 Moroccan accessions (**Figure 1a, Table S1**). Analysis of the population structure using principal component clustering analyses (PCA) (**Figure 1b)** reveals distinct clustering of Santo Antão and Fogo populations relative to the Moroccan population. These findings are consistent with previous results, which indicated that the two islands diverged without any subsequent admixture between them (Tergemina *et al*., 2022; Fulgione *et al*., 2022). A comparison of climate distributions between CVI and Moroccan *A. thaliana* collection sites shows that annual accumulated rainfall amounts are significantly lower in the CVI range relative to the Moroccan range (**Figure 1c**). Additionally, water vapor pressure is higher in CVI compared to Moroccan *A. thaliana* sites (**Figure 1d**). These differences highlight what is likely an important selection pressure in CVI: limited and unpredictable rainfall, with the most reliable moisture deriving from elevated humidity.

**Figure 1.**
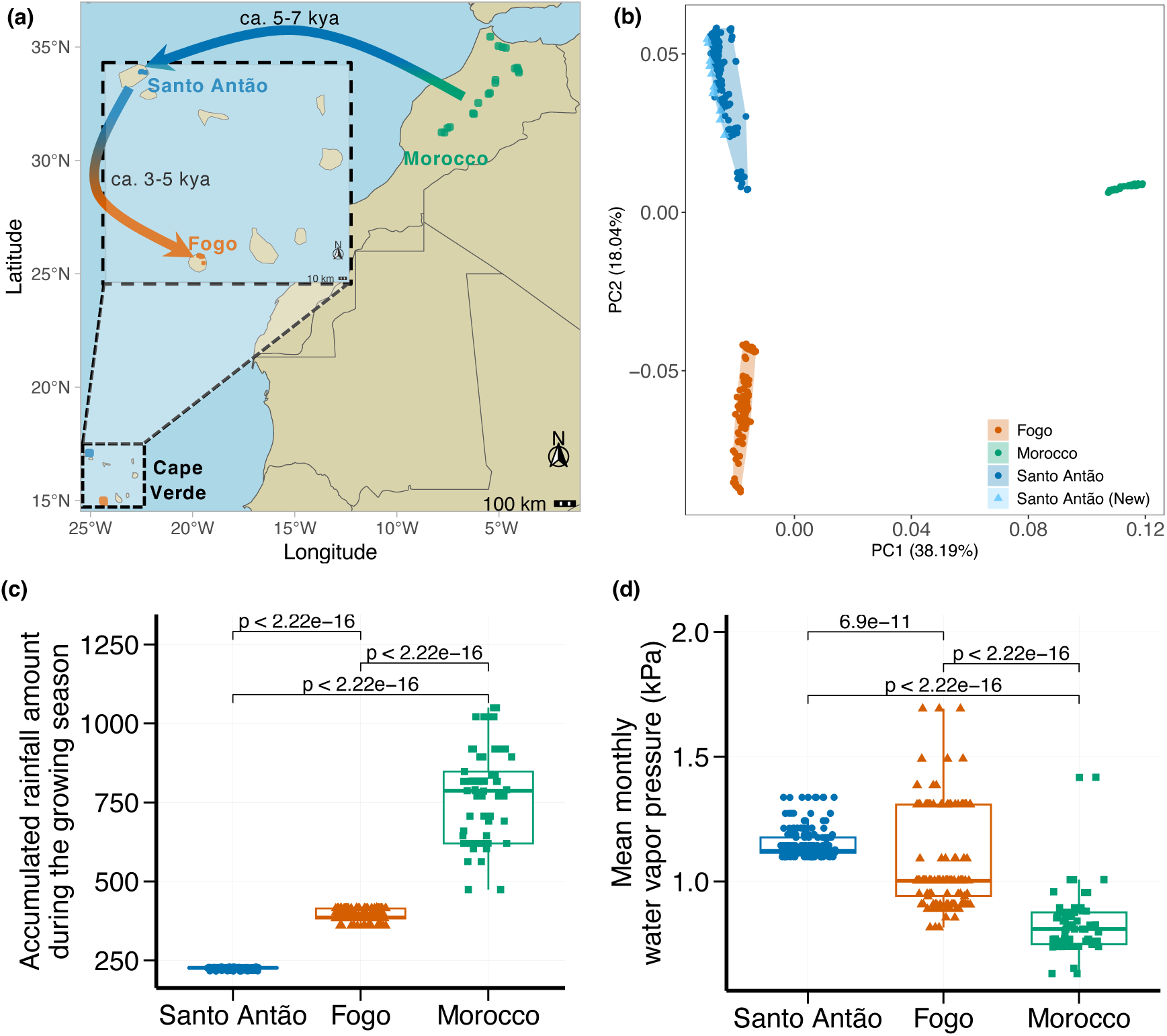
Geographical distribution and genetic structure of the Cape Verde Islands (CVI) and Morocco. (a) Geographic locations populations of *A. thaliana* in CVI (Santo Antão and Fogo) and the outgroup panel of Moroccan *A. thaliana* accessions. (b) PCA of genome-wide SNPs between CVI (Santo Antão and Fogo) and Morocco *A. thaliana* populations. Climate of (c) accumulated rainfall during growing season (kg m^-2^ gsl^-1^), and (d) mean monthly water vapor pressure (KPa) at collection sites in Santo Antão and Fogo relative to Moroccan sites. The line in the center of the boxplots represents the median, the box edges represent the 25th and 75th percentiles (lower and upper bound, respectively), and the whiskers represent 95% CI. The *P*-values for the Mann-Whitney-Wilcoxon test are shown.

### Variation in drought response within CVI and compared to Morocco

We investigated variation in a range of traits (**Table 1**) and their response to moderate drought stress in the CVI islands and the nearest outgroup *A. thaliana* population from Morocco. Traits were assayed under tightly controlled conditions utilizing the Phenoscope, a high-throughput phenotyping platform (Tisné *et al*., 2013). In this platform, pots are automatically and constantly rotated to eliminate the effects of micro-environmental heterogeneity and watering is adjusted multiple times daily to maintain highly controlled soil water content, enabling experiments that would be impractical using manual methods (Tisné *et al*., 2013).

**Table 1.** Key to phenotypic traits.

| Trait ID | Trait description | Unit |
| --- | --- | --- |
| **WUE** | Water Use Efficiency | ‰ |
| **PRA** | Projected Rosette Area | cm<sup>2</sup> |
| **Compactness** | CompactnessPC - ratio between PRA and its convex hull area | NA |
| **RER** | Relative rosette Expansion Rate | day<sup>-1</sup> |
| **Hue** | Rosette average color Hue | % |
| **Saturation** | Rosette average color Saturation | % |
| **SD** | Stomatal Density | 1 mm<sup>-2</sup> |
| **SPW** | Stomatal Pore Width | mm |

We conducted ANOVAs to assess differences between populations in average traits and in the response to drought. Both CVI populations exhibited lower WUE, smaller rosette area, reduced compactness, higher relative expansion rate and lower hue than the Moroccan population (**Figure 2, Table S2**). All traits except Hue shifted significantly overall in response to drought, highlighting phenotypic plasticity in CVI populations under water-limited conditions. We found that drought had a notable impact on leaf growth (**Figure S1**), stomata-related traits, and leaf color-related traits in Santo Antão and Fogo populations compared to Moroccan *A. thaliana* populations (**Figure 2**). Under water deficit (WD) conditions, CVI plants exhibited reduced rosette size, diminished growth rates, and changes in leaf color-related traits compared to well-watered (WW) conditions (**Figure 2a-f**). WUE was significantly lower in the Santo Antão *A. thaliana* population compared to the Moroccan population under both WW conditions and drought conditions (**Figure 2a**). WUE in Fogo was higher than in Santo Antão but lower than in the Moroccan population. These findings are consistent with adaptive responses to localized water regimes, suggesting a complex interplay of genetic and environmental factors driving these differences.

**Figure 2.**
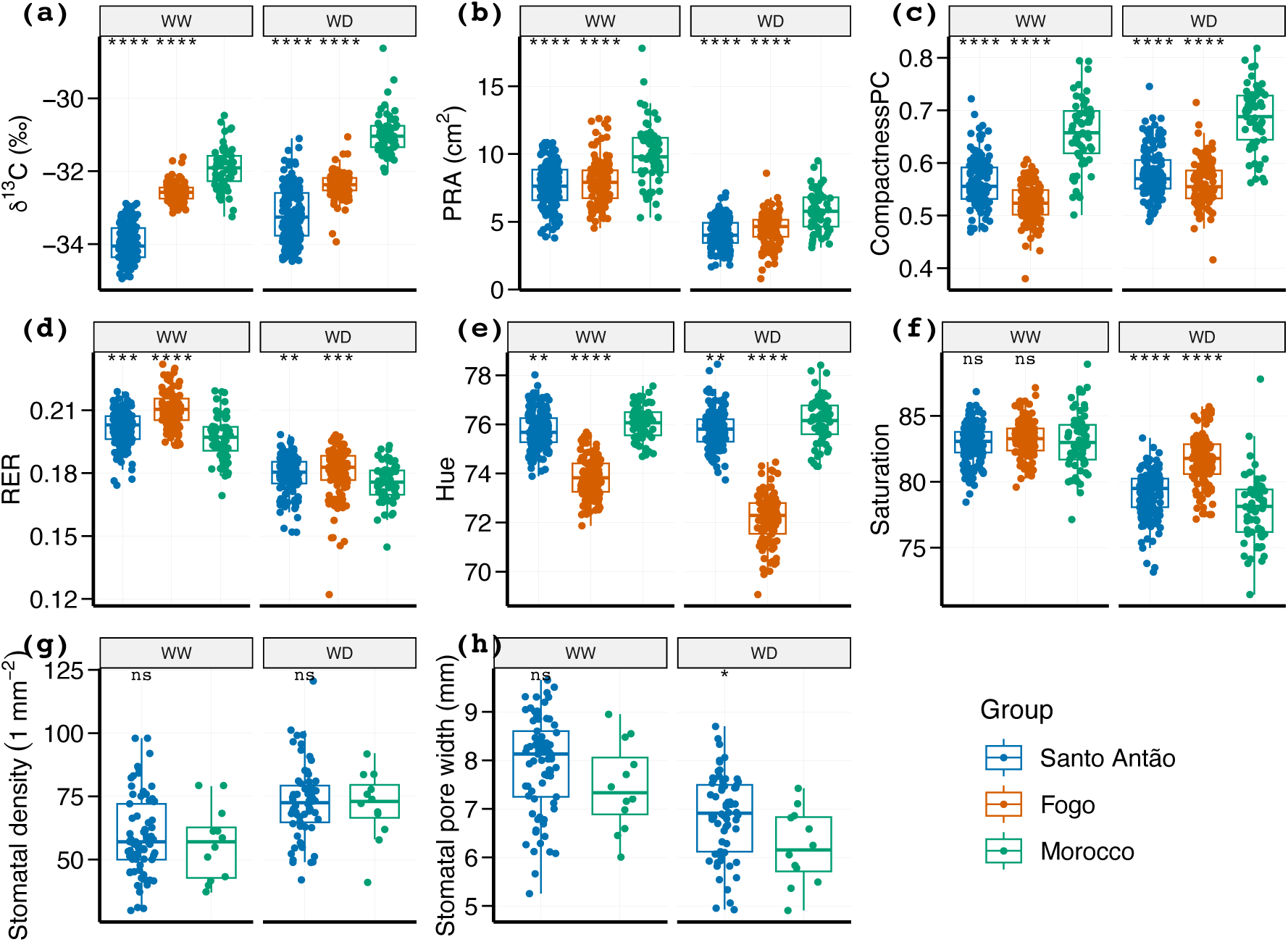
Phenotypic variation in morphological and physiological traits for Santo Antão and Fogo *A. thaliana* populations relative to Moroccan population in well-watered (WW) and water deficit (WD) conditions. The line in the center of the boxplots represents the median, the box edges represent the 25^th^ and 75^th^ percentiles (lower and upper bound, respectively), and the whiskers represent the 95% CI. (a) Water use efficiency (WUE) is measured as carbon isotope discrimination (δ^13^C), and the carbon isotope ratio is expressed per mil, ‰. (b) Projected rosette area (PRA). (c) CompactnessPC (ratio of PRA (P) to surface of convex hull area (C)). (d) Relative expansion rate (RER), the relative growth rate of PRA; integrated over the 18-31 Days After Sowing (DAS) time window. (e-f) Hue is the tint of the rosette color, and saturation represents color intensity. (g) Stomatal density (h) and stomatal pore width. Statistical significance was determined through the Mann-Whitney-Wilcoxon test, using the Moroccan population as a reference. Significance levels are denoted by asterisks, with a single asterisk (*) for *p* ≤ 0.05, double asterisks (**) for *p* ≤ 0.01, triple asterisks (***) for *p* ≤ 0.001, and quadruple asterisks (****) for *p* ≤ 0.0001. Non-significant results are indicated by “ns” for *p* > 0.05.

To delve deeper into the physiological mechanisms underlying the lower WUE in Santo Antão, we examined variation in stomatal density and stomatal pore width (SPW) in a subset of the Santo Antão and Moroccan population samples (**Figure 2g-h**). Stomatal density was similar in the two populations in both well-watered and drought conditions and only the Moroccan population exhibited a significant shift in response to drought. Although mean stomatal pore width was higher in Santo Antão relative to Morocco, the difference was not significant and again only the Moroccan population exhibited a significant decrease in response to drought.

We next estimated trait heritability in each population and condition. Broad-sense heritabilities (*H²*), estimated based on repeatability across replicates, indicated moderate to high heritability for most traits across treatment conditions (**Table S3**). These findings highlight the genetic basis of key drought-related traits and reveal substantial variability in their genetic architecture across populations and environmental conditions.

### Average traits and drought response traits

To better understand phenotypic variation across populations and conditions, we examined variation in average trait values and drought responses. Average traits provide insights into baseline performance and constitutive genetic effects, while drought responses capture phenotypic plasticity and condition-specific adaptations to water deficit. This distinction allows us to evaluate not only overall trait variation in separate conditions but also the degree of plasticity expressed in response to drought stress. Analyzing these complementary aspects may enhance our ability to disentangle genetic mechanisms driving phenotypic shifts under contrasting conditions and strengthen the detection of loci involved in adaptive responses.

We next quantified the proportion of variance explained (PVE) by sequence polymorphisms for average traits and drought responses across populations to assess the relative contribution of genetic effects. Results revealed substantial variation in the PVE across traits and populations (**Figure 3**). Average traits consistently exhibited high PVE (**Figure 3**), particularly compactness in Morocco (0.84, 95% CI 0.57–0.97) and WUE in Santo Antão (0.77, 95% CI 0.67–0.85) and Fogo (0.74, 95% CI 0.52–0.92), highlighting strong genetic effects segregating within these populations. Drought responses showed moderate PVE, reflecting greater plasticity (**Figure 3**). Notably, hue exhibited significant PVE in Santo Antão (0.44, 95% CI 0.26–0.63) and Fogo (0.45, 95% CI 0.25–0.68), while Morocco’s hue trait had the highest PVE (0.66, 95% CI 0.34–0.88) under drought. Overall, these findings demonstrate that while average traits are largely driven by genetic effects, drought responses also exhibit measurable genetic variation, albeit shaped by greater plasticity. This dual approach of examining baseline performance and plastic responses provides a framework for identifying genetic loci underpinning drought adaptation.

**Figure 3.**
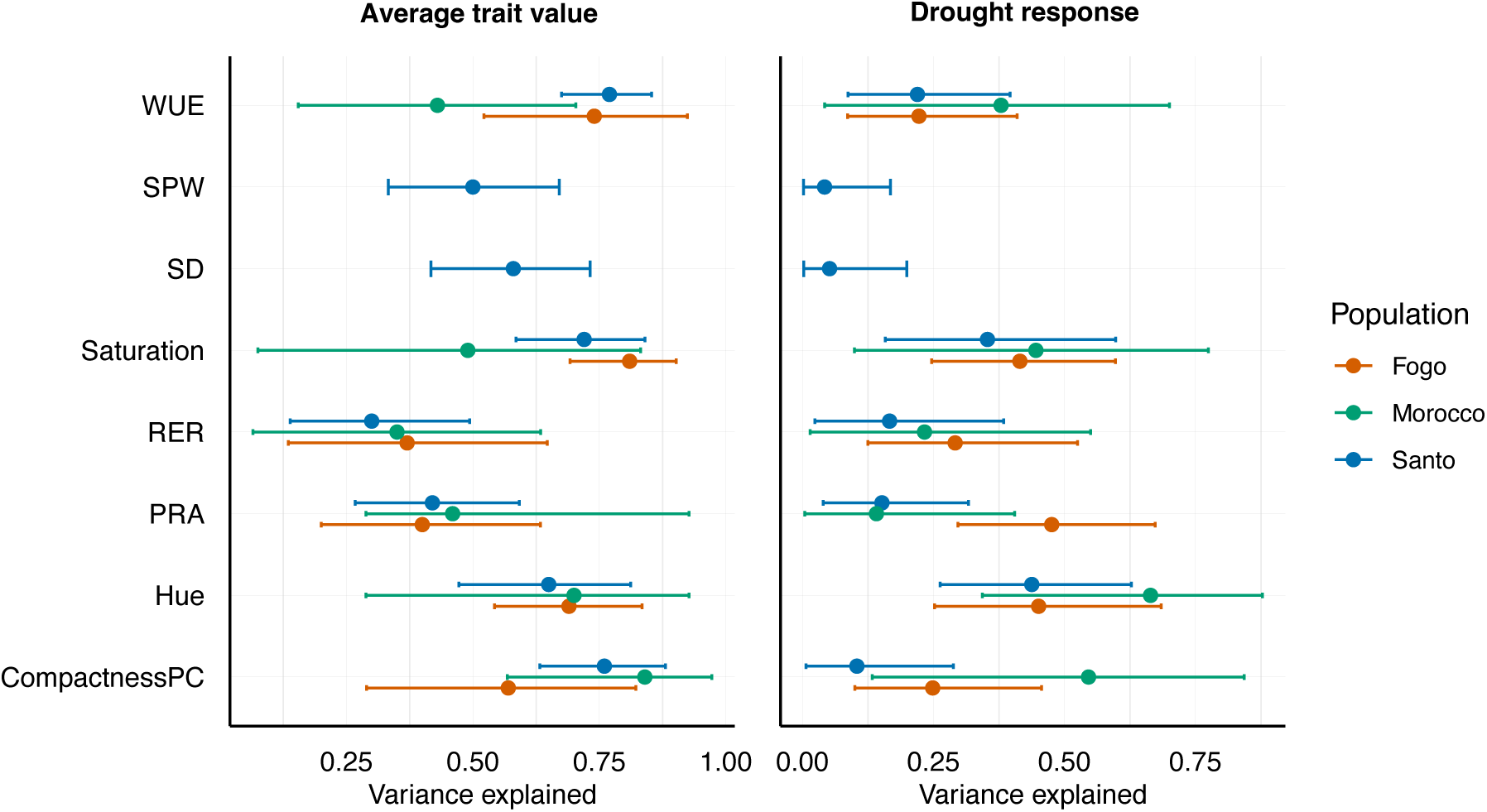
Dot chart shows estimates and confidence intervals for the proportion of variance explained by sequence polymorphisms in the average trait (left) and the drought response (right) in Santo Antão, Fogo, and Morocco populations. PRA, projected rosette area; WUE, water use efficiency; SD, stomatal density; SPW, stomatal pore width; RER, relative expansion rate.

### Phenotypic and genetic correlations between drought-related traits

A shared genetic basis of traits, i.e., pleiotropy, could act to restrict or promote adaptation. To investigate the extent of correlations between pairs of traits and evidence for a shared genetic basis, we estimated genetic correlations for both average traits and drought responses (**Figure 4**, **Figure S2**) using linear mixed models (LMMs), which explicitly control for population structure. Similarly, phenotypic correlations were assessed after accounting for population structure to ensure consistency and comparability between phenotypic and genetic patterns.

**Figure 4.**
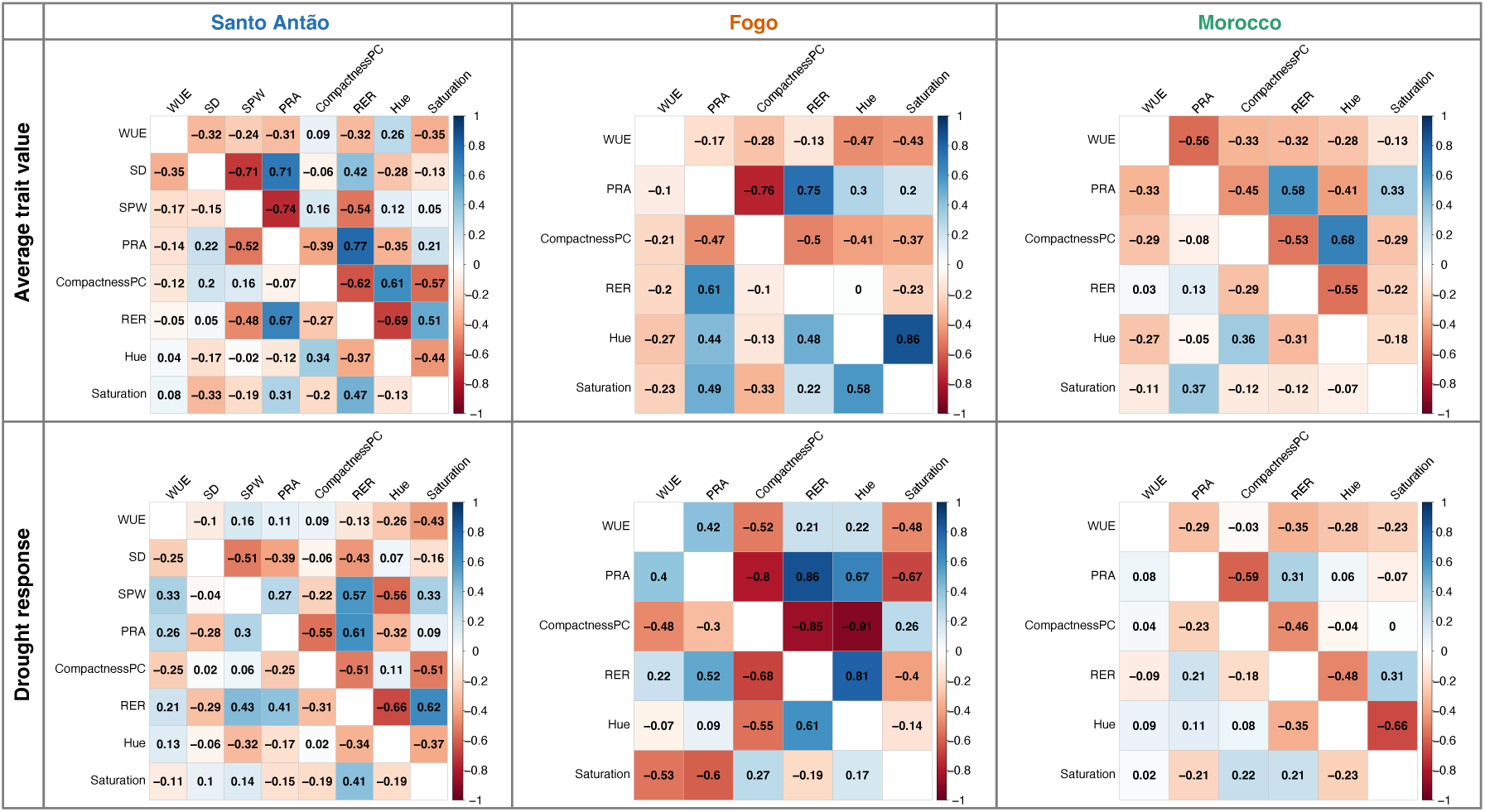
Pairwise Pearson phenotypic (upper right triangle) and genetic (lower left triangle) correlation of the average trait value (upper panel) and the drought response (lower panel) in Santo Antão, Fogo, and Morocco populations. The color spectrum, dark red to dark blue, represents highly negative to highly positive correlations, and the number represents the correlation values.

Overall, we found significant congruence between phenotypic and genetic correlations across drought-related traits in Santo Antão, Fogo, and Morocco (**Figure 4**, **Figure S2**). Similarity between phenotypic and genotypic correlations was especially strong for the traits PRA, RER, and WUE, implying that in these cases, there is likely a strong shared genetic foundation for these drought-related traits. Such genetic-phenotypic correlations are indicative of a complex adaptive mechanism whereby plants fine-tune their growth patterns and WUE in response to drought, governed by underlying pleiotropic functions. This complexity underscores the biological importance of understanding the genetic basis of phenotypic traits, particularly in the context of environmental adaptations.

### Genetic basis of drought response in natural populations

To identify specific loci underlying average traits and drought-response traits, we conducted genome-wide association studies (GWAS). For this, we used an univariate LMM implemented in GEMMA (Zhou & Stephens, 2012), adjusting for population structure by incorporating a kinship matrix as a random effect. We performed GWAS for each trait in each of the two island populations (Santo Antão and Fogo) as well as in the Moroccan outgroup population. After initial mapping using the LMM, we used the local score method (Bonhomme *et al*., 2019) to improve precision and detection power. The local score statistic is calculated by integrating association scores over genomic regions and finding the maximum of a Lindley process over the sequence of these scores (Bonhomme *et al*., 2019). In the Santo Antão population, where we had population-level data for the stomatal traits WUE, SD, and SPW, we also applied a multivariate LMM (mvLMM) approach to examine these as a group (Zhou & Stephens, 2014) because these traits are related and may potentially have a shared genetic basis. By employing these strategies, we could identify significant associations across the genome. **Figure 5** highlights the most relevant candidate genes associated with average traits and drought response traits in the three populations. Individual GWAS results are provided in **Figure S3-9** and **Table S4-S6**.

**Figure 5.**
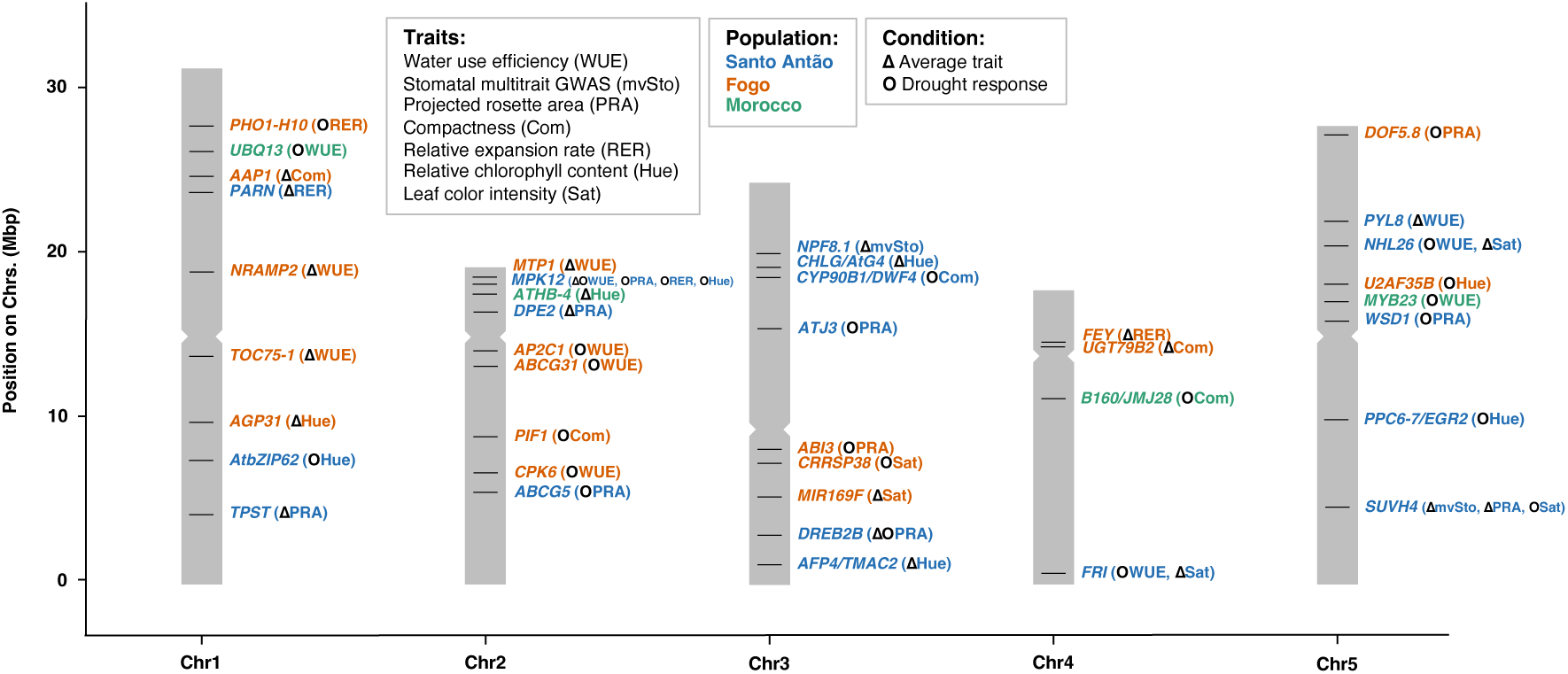
Genomic associations across eight traits in response to drought identified by GWAS. Associations shown include SNPs linked to 39 candidate genes, selected based on their harmonic mean rank and functional relevance to drought-related traits (see Materials and Methods). Traits (see abbreviations) were analyzed across Santo Antão, Fogo, and Morocco populations (distinguished by color) and two treatment conditions (represented by different shapes). Detailed annotations of all SNPs are available in **Table S4-S6**.

GWAS identified several potential candidate genes associated with rosette growth under drought stress, including those shown to be involved in auxin signaling (*TPST*), starch turnover (*DPE2*), cuticle formation (*ABCG5*), wax ester accumulation (*WSD1-like*), and stress-responsive signaling (*MPK12*, *DREB2B*, *ATJ3*). Additional published evidence supports the role of these loci in drought response. *TPST* mutants exhibit a small rosette phenotype, early senescence, and a reduced number of flowers and siliques (Komori *et al*., 2009). Similarly, *DPE2* mutants show reduced leaf and rosette area under drought conditions with respect to wild-type, and they also display altered transpiration phenotypes, highlighting the gene’s potential role in coordinating growth and water regulation under stress (Massonnet *et al*., 2015; Westgeest *et al*., 2023). ABCG5 is crucial for cuticle formation, which helps in retaining water and supporting plant growth under stress (Lee *et al*., 2021). Additionally, *WSD1* plays a role in accumulating wax esters, which are vital for reducing water loss (Patwari *et al*., 2019). *DREB2B*, which is induced by drought stress, regulates the transcription of drought-responsive genes through binding to the DRE/CRT sequence (Liu *et al*., 2019). This gene is known for its role in enhancing drought tolerance by regulating stress-responsive gene expression (Nakashima *et al*., 2000; Sakuma *et al*., 2006). *ATJ3*, a member of the HSP40 family, contributes to heat stress tolerance by interacting with HSP70-4 and participating in heat stress granules (Wang *et al*., 2021). These findings highlight potential interplay between metabolic pathways and drought adaptation.

Loci identified in the Fogo population exhibited distinct genetic associations compared to those in the Santo Antão population, which aligns with their early separation and lack of shared genetic variation. In Fogo, loci associated with rosette growth under drought stress included *DOF5.8, UGT79B2, PIF1, FEY, PHO1-H10,* and *AAP1*. *DOF5.8*, a transcription factor, regulates the *ANAC069* gene, which is involved in integrating auxin and salt signals for seed germination (He *et al*., 2015). *UGT79B2*, induced by various abiotic stresses, enhances plant tolerance by modulating anthocyanin accumulation (Li *et al*., 2017). *PIF1* influences ABA biosynthesis and signaling component genes (Oh *et al*., 2007), while *FEY* is implicated in leaf positioning and meristem maintenance (Callos *et al*., 1994). *PHO1-H10*, involved in phosphate homeostasis, is upregulated by ABA, suggesting its role in drought response through modulating phosphate availability (Ribot *et al*., 2008). *AAP1*, whose expression is induced by ABA, may play a crucial role in amino acid transport under drought stress (Wang *et al*., 2017). Despite differences in specific genes and variants identified, these findings suggest that similar genetic mechanisms may underlie drought responses in both Santo Antão and Fogo populations.

We also investigated leaf color traits, hue and saturation, which represent complex proxies for pigments content (including chlorophyll). Hue values, which have been found to correlate with chlorophyll content under certain conditions and genotypes, may provide indirect insights into photosystem II photochemical yield (Faragó *et al*., 2018). Although previous research has indicated that the saturation parameter was less important than hue values in specific experiments (De Vylder *et al*., 2012), we discovered an association between average saturation and the *FRIGIDA* (*FRI*) gene, which influences various traits, including flowering and WUE (Lovell *et al*., 2013). This suggests that *FRI* may have pleiotropic effects on drought adaptation, influencing both leaf color and WUE.

A combined gene ontology (GO) enrichment analysis across the eight drought response traits revealed significant enrichment for gene sets involved in drought response, providing evidence that the GWAS was detecting true associations (**Figure 6, Table S7**). We found that terms related to ‘response to abscisic acid’ were overrepresented for the candidate genes.

**Figure 6.**
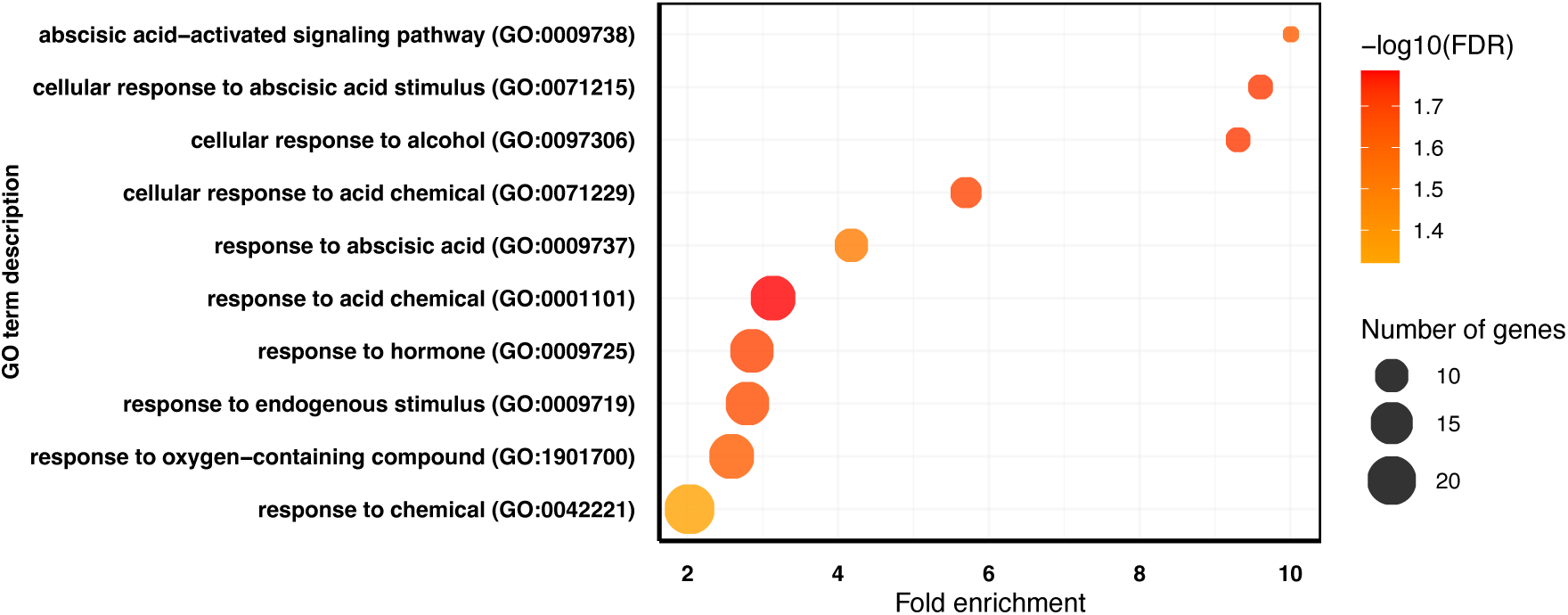
GO term enrichment analysis for the candidate genes identified across eight traits in response to drought. The size of the circles indicates the number of genes and the circle color the corresponding *P*-value for enrichment after FDR correction.

This was driven by genes such as *ABI1* and *PYL8* (**Figure 6**), known ABA receptor components (Belda-Palazon *et al*., 2018), and *MPK12*, which is implicated in ABA signal transduction, *VQ18*, and *DREB2B*, known for its involvement in drought-responsive pathways (Liu *et al*., 2019).

Co-localization of GWAS peaks across traits indicates potential pleiotropic effects of some of the identified loci, including *MPK12*, *FRI*, *NHL26*, and *SUVH4*. We found that *MPK12* not only associates with WUE but also with drought response of PRA, RER, and Hue, reflecting the strong genetic correlations observed among these traits. This suggests that *MPK12* may act as a central regulator, mediating coordinated responses to drought stress across physiological and morphological traits. Furthermore, a locus that includes a premature truncation of *FRI*, a gene traditionally known for its role in flowering time regulation, was associated with WUE and leaf color intensity (saturation) (**Figure 5, Figure S9a**), indicating a broader adaptive response to drought. Additionally, *SUVH4*, an epigenetic regulator via H3K9 methylation, was associated with stomatal traits, PRA and leaf color, suggesting a link between epigenetic modifications and the physical adaptation to drought, providing evidence that epigenetic mechanisms that may modulate drought stress responses. Together, these findings underscore how pleiotropic loci, identified through GWAS and supported by genetic correlations, can shape coordinated phenotypic responses to drought, enabling plants to adapt to water-limited environments.

## Discussion

Here we investigated drought response in *A. thaliana* populations from the Cape Verde Islands (CVI) in comparison to their closest outgroup populations from Morocco. We used a high-throughput phenotyping platform (Phenoscope) to measure key traits under controlled drought conditions and GWAS to connect trait variation with its genetic underpinnings. CVI populations showed significant shifts in the distribution of traits relative to Morocco, including lower water use efficiency (WUE), smaller rosette size, reduced compactness, and higher relative growth rates. Genetic correlations between traits and congruence in GWAS peaks across some trait pairs provided evidence for pleiotropy. Furthermore, we observed an enrichment for genes involved in abscisic acid (ABA)-related pathways, underscoring the pivotal role of ABA in modulating drought related traits and promoting adaptive responses.

In the GWAS analysis, we identified candidate loci involved in ABA signaling and epigenetic processes. The enrichment of candidate genes in the ABA signaling pathway reflects its established role in modulating drought responses, such as stomatal closure, water uptake, and osmotic adjustment (Sharma *et al*., 2011; Tuberosa, 2012). ABA-related pathways are known to influence physiological processes like stomatal regulation and proline metabolism, both critical for plant resilience to drought (Setter, 2012; Bhargava & Sawant, 2013). Our GWAS findings identified *NPF8.1*, a key gene facilitating ABA uptake into guard cells, which modulates stomatal closure (Tal *et al*., 2016; Shimizu *et al*., 2021). This complements prior observations linking ABA signaling to proline accumulation and oxidative stress mitigation (Bhatt *et al*., 2011; Liang *et al*., 2013). Additionally, *SUVH4*, influenced by ABA modulation (Zheng et al., 2012), was implicated in multiple drought-related traits, including stomatal density, leaf color and plant rosette area, suggesting pleiotropic effects. Overall, these results highlight the central role of ABA signaling in coordinating drought-related responses and shaping trait variation across populations, and they reveal the intriguing possibility that epigenetic processes might also contribute to drought response.

The relationship between flowering time and vegetative growth dynamics is well-established (Bac-Molenaar *et al*., 2016; Hanemian *et al*., 2020). Our study suggests that pleiotropy may play a role in drought adaptation. Previous studies have demonstrated that correlated physiological traits, such as WUE and flowering time, can influence adaptive processes (McKay *et al*., 2003; Lovell *et al*., 2013). Our findings complement these studies by identifying genes like *FRIGIDA* (*FRI*), which may link leaf color traits to adaptive processes such as the transition from vegetative growth to flowering.

*FRI*, previously associated with reduced time to flowering time and increased seed production in CVI environmental conditions (Fulgione *et al*., 2022), appears to play a broader role in adaptation to drought environments. The presence of the *FRI* 232X variant in Santo Antão suggests it represents an early step in adaptation to the CVI environment, facilitating a drought-escape strategy by accelerating reproduction before water resources become limiting (Fulgione *et al*., 2022; Neto & Hancock, 2023). Beyond its established role in flowering time regulation, *FRI* was significantly associated with saturation, a trait related to leaf color intensity, which may reflect shifts in resource allocation from vegetative growth to reproductive success, a key feature of drought escape strategies. This observation highlights a potential pleiotropic function of *FRI*, linking WUE and leaf color intensity to reproductive timing and investment. In particular, *FRI*’s association with WUE underscores its role in optimizing carbon assimilation while minimizing water loss, enabling plants to balance rapid growth and reproduction under drought conditions. This suggests that the *FRI* locus may mediate coordinated responses across multiple physiological and developmental pathways, aligning traits critical for survival and reproduction under drought. The pleiotropic effects of *FRI* on traits related to growth, WUE, and reproduction illustrate its integrative role in shaping the escape strategy observed in CVI populations.

Here, we found evidence that *MPK12* plays a role in multiple drought-related traits, including WUE, stomatal traits, growth, and leaf color. Previous studies have established *MPK12* as a key regulator of stomatal responses to ABA, influencing WUE and guard cell behavior, with the CVI allele altering responses to vapor pressure deficits and ABA-induced inhibition of stomatal opening (Juenger *et al*., 2005; Jammes *et al*., 2009; Salam *et al*., 2013; Montillet *et al*., 2013; Des Marais *et al*., 2014; Jakobson *et al*., 2016). In earlier work on the Santo Antão population, a nonsynonymous variant in *MPK12* (G53R) was shown to drive phenotypic shifts towards higher stomatal conductance and lower WUE (Elfarargi *et al*., 2023). These shifts supported a drought-escape strategy in Santo Antão, where plants maximize carbon assimilation through open stomata under conditions of high atmospheric humidity (Elfarargi *et al*., 2023). Incorporating this variant as a covariate in our GWAS model allowed us to uncover additional loci associated with ABA pathways. Notably, *MPK12* was linked to WUE, stomatal traits, growth, and leaf color, indicating its broader pleiotropic role in drought-related adaptation. The ability of populations to adapt rapidly to the extreme climate shifts in CVI likely relies on balancing loci with targeted functions and pleiotropic effects.

This tradeoff facilitates efficient adaptation for managing complex traits under selection. Our findings support this hypothesis, with *MPK12* emerging as a central hub in the genetic network underlying drought response.

Despite genetic and allelic heterogeneity between Santo Antão and Fogo populations, our GWAS results reveal a convergence on shared genetic pathways underlying drought responses. While the specific genes identified varied between populations, both islands exhibited enrichment in pathways related to ABA signaling and growth regulation, suggesting that similar physiological processes are being targeted by natural selection.

Although the genetic components differ, the underlying functional processes, such as regulating WUE, stomatal behavior, and stress signaling, are remarkably consistent. This suggests that these populations are adapting to drought through distinct genetic mechanisms converging on shared pathways. Such insights deepen our understanding of how distinct populations achieve functional convergence despite genetic divergence, highlighting the complex interplay between local adaptation and shared evolutionary pressures.

While this study highlights promising candidate genes and their associations with drought-related traits, further validation is required to establish causality. Functional studies using knockout lines, reciprocal transformations, and gene expression analyses will be essential to confirm these candidate genes as drivers of phenotypic variation. Additionally, integrating co-expression network analysis could unravel the interplay between loci with targeted and pleiotropic effects. Future studies might also extend these findings to crop systems, leveraging the identified genes to improve drought tolerance in agriculturally important species.

This study provides new insights into how plants have evolved to cope with changes in precipitation regimes. It extends our understanding of drought adaptation mechanisms by identifying significant genetic associations linked to key drought-related traits. Further, our findings underscore the potential of pleiotropy to facilitate adaptation. This research paves the way for a deeper understanding of plant responses to abiotic stress and establish a framework for exploring adaptive mechanisms in both natural populations and agricultural systems.

## Materials and methods

### Study populations

In this study, we used a total of 350 accessions of *Arabidopsis thaliana* from the Cape Verde Islands (CVI) and Morocco to investigate genetic and phenotypic variation associated with drought adaptation. Our sampling included 189 accessions (Fulgione *et al*., 2022) along with 15 newly collected accessions from 26 locations in Santo Antão (**Figure 1b**). Additionally, we examined 146 accessions collected across 18 locations in Fogo (Tergemina *et al*., 2022; Fulgione *et al*., 2022) and 61 Moroccan accessions (Durvasula *et al*., 2017) as an outgroup population.

We generated maps to visualize geographic sampling locations using R v.3.4.4 (R Development Core Team, 2008), and employed the *ggmap* (Kahle & Wickham, 2013) and *ggplot2* (Wickham, 2016) libraries for plotting.

### Whole-Genome Sequencing

We sequenced 15 new Santo Antão accessions using Illumina HiSeq3000 machines. Genomic DNA was extracted using the DNeasy Plant Mini kits (Qiagen), fragmented using sonication (Covaris S2), and libraries were prepared with Illumina TruSeq DNA sample prep kits (Illumina), and NEBNext Ultra II FS DNA Library Prep Kit (New England Biolabs).

Libraries were sequenced with 2x 100-150 bp paired-end reads. We assessed DNA quality and quantity via capillary electrophoresis (TapeStation, Agilent Technologies) and fluorometry (Qubit and Nanodrop, Thermo Fisher Scientific).

### Variant identification and genotyping

Here, we called variants from the newly sequenced 15 Santo Antão accessions together with the previously released whole-genome short-read data for 189 Santo Antão accessions (ENA: PRJEB39079 (ERP122550)) (Fulgione *et al*., 2022). We aligned the raw Illumina sequence for the whole data to the *Arabidopsis* TAIR10 reference genome. Further, we used SHORE (Ossowski *et al*., 2008) to identify SNP variants following a pipeline described here (https://github.com/HancockLab/SNP_calling_Arabidopsis). Furthermore, the obtained variant call format (VCF) file was filtered to minimize variant calling bias and to retain only high-quality variants: (1) retain only bi-allelic variants; (2) convert heterozygous sites to missing data to mask possible false positives; (3) retain variants with coverage greater than 3 and base quality greater than 25.

### Population structure analysis

We analyzed population structure by first pruning SNPs for linkage disequilibrium using PLINK v1.9 (Purcell *et al*., 2007). SNPs with pairwise correlation coefficients (*r²*) > 0.1 were removed within 50 Kb sliding windows with a 10 bp step size. Additionally, we excluded variants with missing genotype data by setting the --geno parameter to 0. We conducted a principal component analysis (PCA) to summarize genetic variation patterns among populations using the --pca function in PLINK v1.9.

### Phenoscope drought experiment and phenotyping

We conducted phenotypic measurements using the high-throughput Phenoscope platform (https://phenoscope.versailles.inra.fr/) as previously described (Tisné *et al*., 2013). A total of 359 accessions of *Arabidopsis thaliana* were included: 152 from Santo Antão, 146 from Fogo (Tergemina *et al*., 2022; Fulgione *et al*., 2022) and 61 from Morocco (Brennan *et al*., 2014). Plants were grown under controlled environmental conditions (8-h day/16-h night, 21°C day/17°C night, 65% relative humidity, and 230 µmol m^-2^ s^-1^ light intensity). Two watering conditions were implemented: a well-watered (WW) condition where pots were maintained at 60% of maximum soil water content (SWC; 4.6 g H₂O/g dry soil), and a water-deficit (WD) condition where pots were maintained at 25% SWC (1.4 g H₂O/g dry soil). The Phenoscope setup, water-use efficiency (WUE) measurements, and sample collection protocols can be found here (Elfarargi *et al*., 2023). The image-based traits were defined as follows: projected rosette area (PRA), hue and saturation (converted from rosette RGB color components), compactnessPC (ratio of PRA (P) to convex hull area (C)) were extracted at final timepoint (31 Days After Sowing (DAS)). Relative Expansion Rate (RER) was integrated from PRA over time window 18 DAS to 31 DAS.

In this study, we measured further leaf stomatal density and stomatal pore width in the Santo Antão population. We adapted a previously published protocol, “Microscopy-Based Stomata Analyses” (Eisele *et al*., 2016), using a small piece of Scotch tape. The tape was applied firmly onto the abaxial surface of a detached fully developed leaf per accession and per treatment of each genotype. Then, we gently peeled off the tape from the leaf surface, not tearing it. We transferred the tape immediately to a petri dish containing 4% formaldehyde solution for 10-20 minutes at room temperature to fix the status of stomatal guard cells and prevent their responses to subsequent signals. Next, we put the samples (tapes with epidermal peels) into a clean tissue and added 50 µl of 1 mg/ml propidium iodide solution to each sample and incubated them for 5-10 minutes in the dark. Then we placed the tape on a microscope slide and gently pressed it to remove any air bubbles. As a final step, we examined for stomatal densities and stomatal apertures using confocal images of epidermal peels taken with a confocal microscope (LSM 700, ZEISS). The images were captured and analyzed using ImageJ software (https://imagej.nih.gov/ij/). The measurements were performed across several days by the end of the Phenoscope experiment (29-32 DAS) around mid-day.

### Phenotype data analysis

We assessed differences in phenotype distributions using both parametric and non-parametric tests. Wilcoxon rank sum tests were performed with the *wilcox.test* function from the ‘*ggpubr*’ package (Kassambara, 2020) to evaluate pairwise comparisons. To analyze the fixed effects of treatment, geographic region, and their interaction on phenotypic traits, we employed linear mixed models implemented in the ‘*lme4*’ package in R (Bates *et al*., 2014). For each phenotype, we fit the following model:

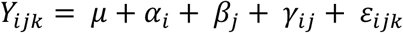

Where *Y_ijk_* represents the phenotypic value, μ is the overall mean, α*_i_* is the effect of treatment, β*_j_* is the effect of geographic region, *γ_ij_* is the interaction between treatment and region, and ε*_ijk_* is the residuals.

Correlation analyses were conducted using the *cor.test* function in R to evaluate relationships between phenotypes in the average trait value and in response to drought. The significance of the correlations was evaluated with the t-test implemented in the *cor.test* function.

### Genome-wide association study

- **Univariate and Multivariate GWAS**

We performed genome-wide association analyses to identify genetic variants linked to drought-related traits. First, we filtered the SNP data to retain only biallelic variants. Variants were filtered based on read coverage (DP ≥ 3) and genotype quality (GQ ≥ 25), followed by applying a minor allele frequency (MAF ≥ 5%) cutoff to exclude rare variants. Subsequently, we carried out the association analyses between genomic variants and studied traits using the univariate linear mixed model implemented in GEMMA (Zhou & Stephens, 2012), separately for well-watered and drought conditions as well as the average for each trait across both conditions and the drought response (difference between conditions: WW-WD). For stomatal traits, we utilized a multivariate linear mixed model (mvLMM) in GEMMA (Zhou & Stephens, 2014), which jointly models the relationships between the traits.

Following Shim *et al*. 2015 (Shim *et al*., 2015), we calculated the proportion of variance explained (PVE) by each SNP based on GEMMA outputs. The PVE was computed using the following equation:

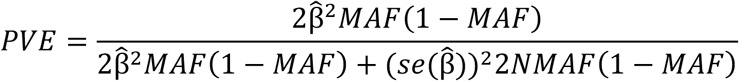

where 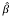 is the effect size estimate, 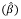 is the standard error of effect size for the variant, *MAF* is the minor allele frequency for the variant, and *N* is the sample size.

Genetic correlations were calculated in R using the *cor.test* function to assess the correlations between the effect size estimate 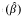 for each trait in the average trait value and in response to drought. The significance of the correlations was evaluated with the t-test implemented in the *cor.test* function.

Next, we applied a local score approach to the output of *p*-values provided by LMM and mvLMM in GEMMA. This local score approach, which takes LD into account when estimating associations, increases the power of detecting significant genomic regions associated with trait variation and is subsequently converted into scores (Bonhomme *et al*., 2019). Scripts used to calculate the Lindley scores are available here (https://forge-dga.jouy.inra.fr/documents/940).

- Inference of genetic architecture

To infer the genetic architecture of drought-related traits, we used the Bayesian Sparse Linear Mixed Model (BSLMM) implemented in GEMMA (Zhou & Stephens, 2012). This polygenic model accommodates both large-and small-effect loci and accounts for relatedness among individuals by including a genomic kinship matrix as a random effect. Furthermore, the approach accounts for the LD between variants by inferring locus effect sizes (β) while controlling for other variants included in the model. Using this approach, we modeled two effect hyperparameters: a basal effect (β), which captures small-effect loci contributing to the studied trait, and an additional effect (γ), which captures a subset of loci with the most potent effects. To estimate the effects of all variants, the sparse effect size for each locus was calculated by multiplying (β) by (γ). We used the 1% variants (> 99% quantile) distribution of locus effects to identify which genes with the highest sparse effects on the studied trait.

- Candidate loci identification and GO enrichment analysis

To identify candidate loci associated with drought-related traits, we integrated results from the GWAS models described above. For each trait and condition within each population, GWAS results were ranked according to their respective association metrics. These ranks were subsequently transformed using the harmonic mean rank, ensuring contributions from all GWAS models were equally weighted. SNP variants within the top 5% of the harmonic rank distribution were selected as significant candidates. From this subset, we highlighted the most relevant candidate genes associated with average traits and drought response traits across populations (**Table S4-S6**). For downstream analyses, gene annotations corresponding to the top-ranked SNPs were retrieved using the TAIR10 GFF3 gene annotation file processed with SNPEff (Cingolani *et al*., 2012). Gene ontology (GO) enrichment analysis was performed to determine if specific biological processes were overrepresented among the candidate loci. GO enrichment analysis was conducted using the ShinyGO tool (Ge *et al*., 2020).

## Acknowledgments

We thank the members of the Hancock Lab provided helpful discussion and feedback. We are grateful to Ångela Moreno and Samuel Gomes at INIDA, as well as the Parque Natural do Santo Antão, for helpful advice and interactions during the course of this research. Philipp Westhoff and Maria Graf provided technical assistance with the δ^13^C analysis. Lisa Schumacher helped with the screening and imaging of stomatal traits. This work was supported by Max Planck Society Funding and European Research Council (ERC) CVI_ADAPT 638810 and the Deutsche Forschungsgemeinschaft (DFG, German Research Foundation) – Project-ID 456082119 – TRR 341/1 to AMH. The International Max Planck Research School (IMPRS) Program “Understanding Complex Plant Traits using Computational and Evolutionary Approaches” partially supported AFE. This work has benefited from the support of IJPB’s Plant Observatory platform PO-Pheno. The IJPB benefits from the support of Saclay Plant Sciences-SPS (ANR-17-EUR-0007). The CEPLAS Metabolomics and Metabolism Laboratory is supported by a grant of the Deutsche Forschungsgemeinschaft (DFG, German Research Foundation) under Germanýs Excellence Strategy – EXC-2048/1 – project ID 390686111.

## Author contributions

AFE and AMH conceived and designed the study. OL provided expertise for designing the drought measurement (Phenoscope) experiment. AFE, OL, and EG conducted the Phenoscope drought experiment and collected samples for δ13C analysis. NK, APMW, and members of APMW’s lab performed δ13C measurements. AFE prepared data, conducted statistical analyses, and created figures. AFE and AMH analyzed and interpreted the results. AFE, AMH, and HD collected plant samples. AFE and AMH wrote the manuscript, with input from all authors. All authors reviewed and approved the final manuscript.

## Data availability

All scripts used for data processing and analysis are available at https://github.com/HancockLab/CVI-DroughtAdaptation-MultiTraitGWAS.

## Supplementary figures

**Figure S1.**
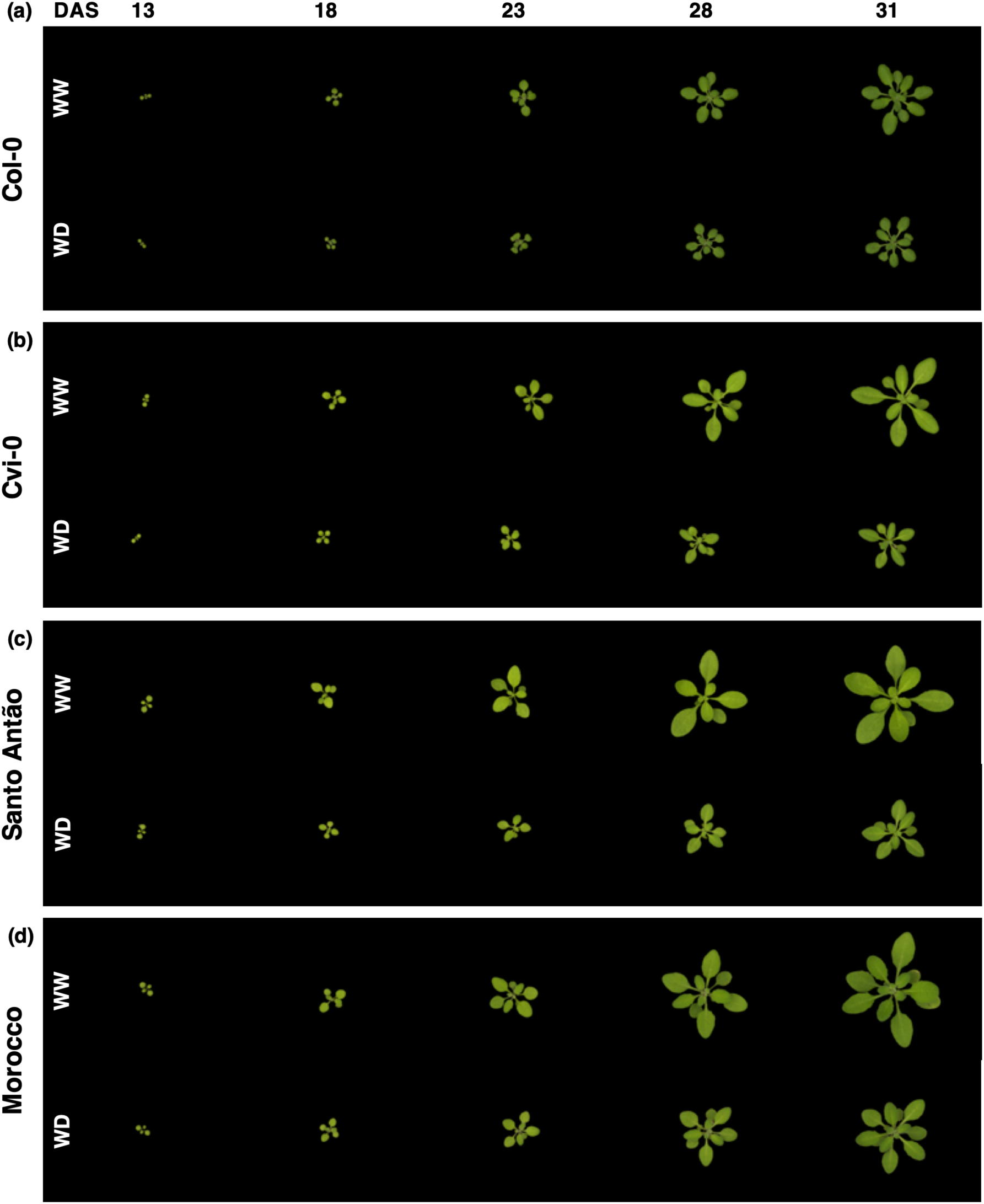
Images of representative accessions on time-points (13, 18, 23, 28, and 31 Days After Sowing (DAS)) of Phenoscope drought experiment in well-watered (WW) and water deficit (WD) conditions. (a) *Col-0*, (b) *Cvi-0*, (c) *S15-3* as a representative accession from Santo Antão, and (d) *Elh-2* as a representative accession from Morocco.

**Figure S2.**
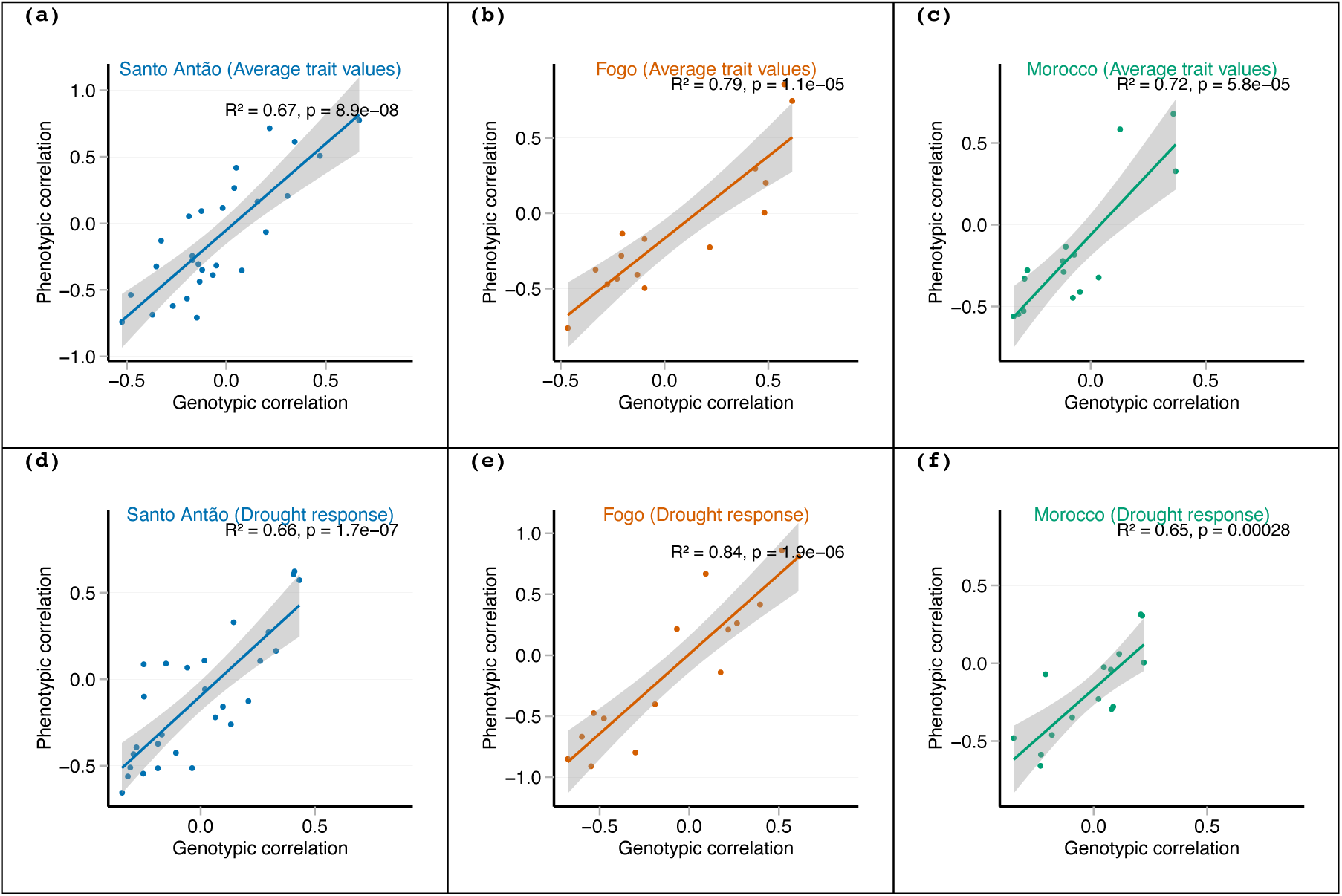
Genetic versus phenotypic correlations across trait pairs for the average trait values in Santo Antão (a), Fogo (b), and Morocco (c), and for the drought response in Santo Antão (d), Fogo (e), and Morocco (f).

**Figure S3.**
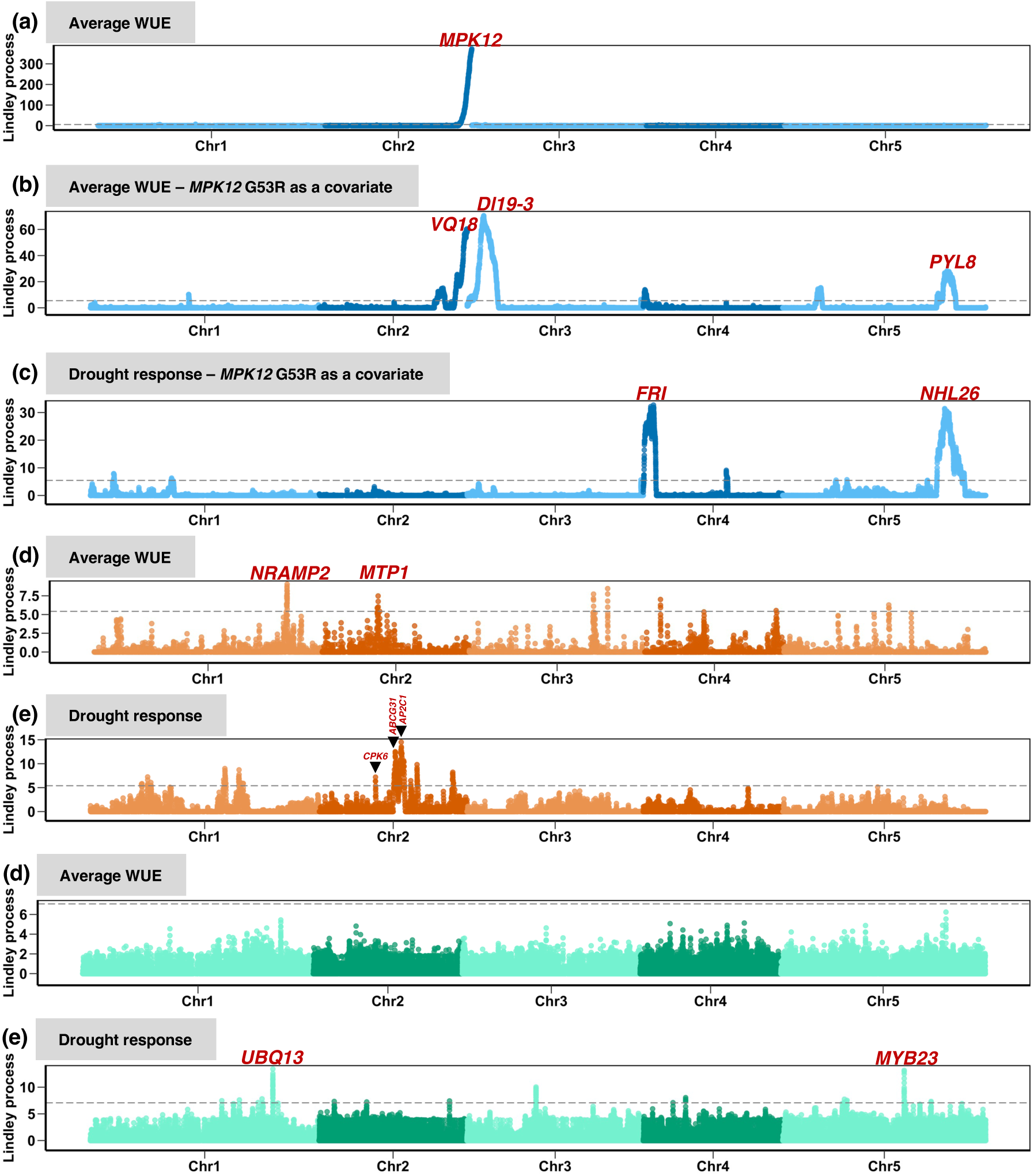
Genomic associations of water use efficiency (WUE) in response to drought in Santo Antão (blue), Fogo (orange), and Morocco (green) populations using LMM in GEMMA followed by the local score approach. The y-axis represents the Lindley process score from the local score approach. The dashed line in plots corresponds to a genome-wide Bonferroni significance.

**Figure S4.**
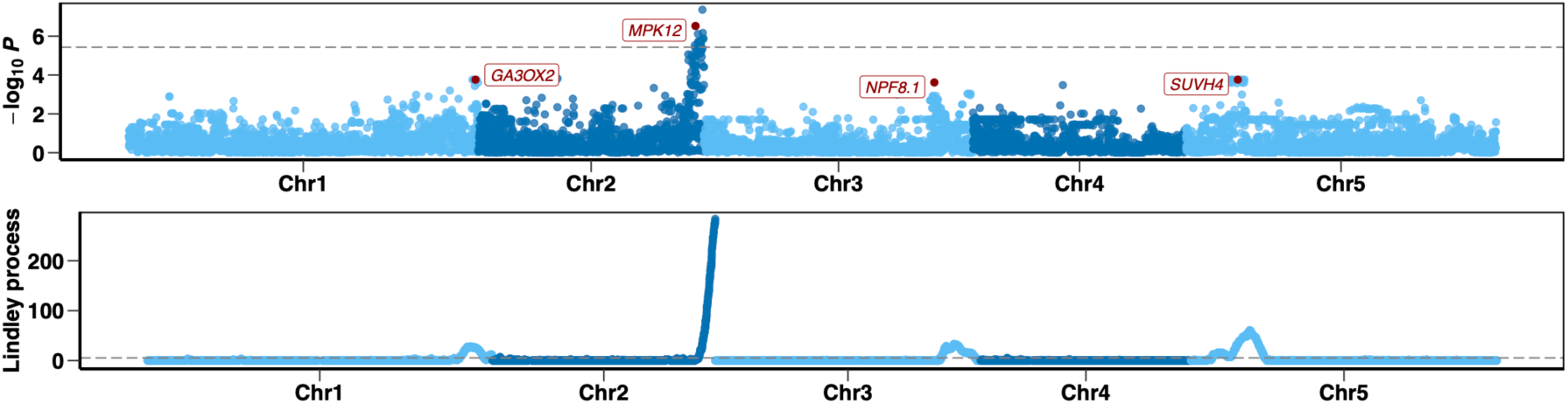
Genomic associations in Santo Antão for stomatal-related traits (WUE, SD, and SPW) using the multivariate model in GEMMA (upper) followed by the local score approach (bottom). The dashed line in plots corresponds to a genome-wide Bonferroni significance.

**Figure S5.**
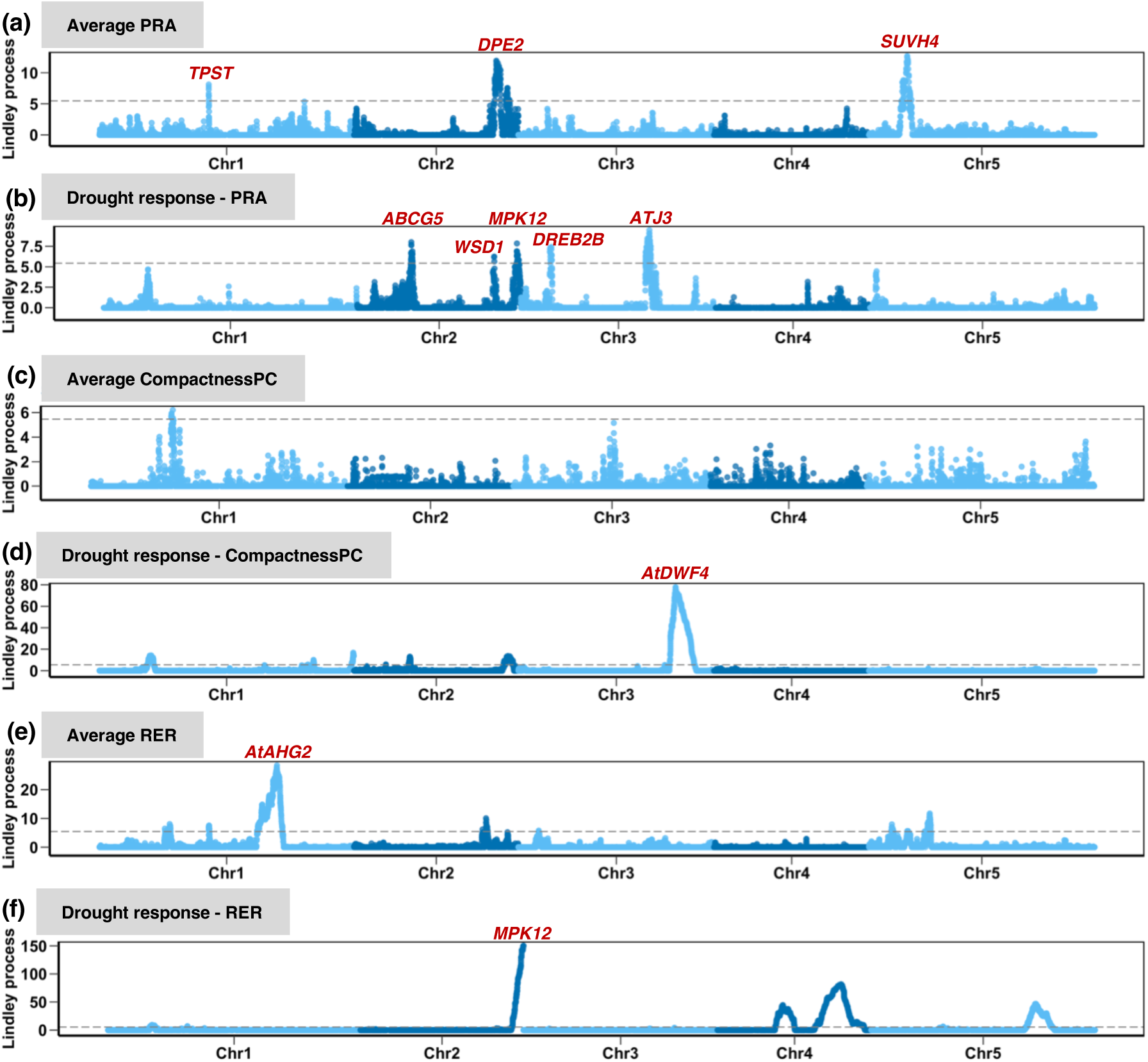
Genomic associations of rosette growth-related traits (PRA, compactnessPC, and RER) in Santo Antão using LMM in GEMMA followed by the local score approach. The x-axis and y-axis represent the chromosomes and Lindley score from the local score approach. The dashed line in plots corresponds to a genome-wide Bonferroni significance.

**Figure S6.**
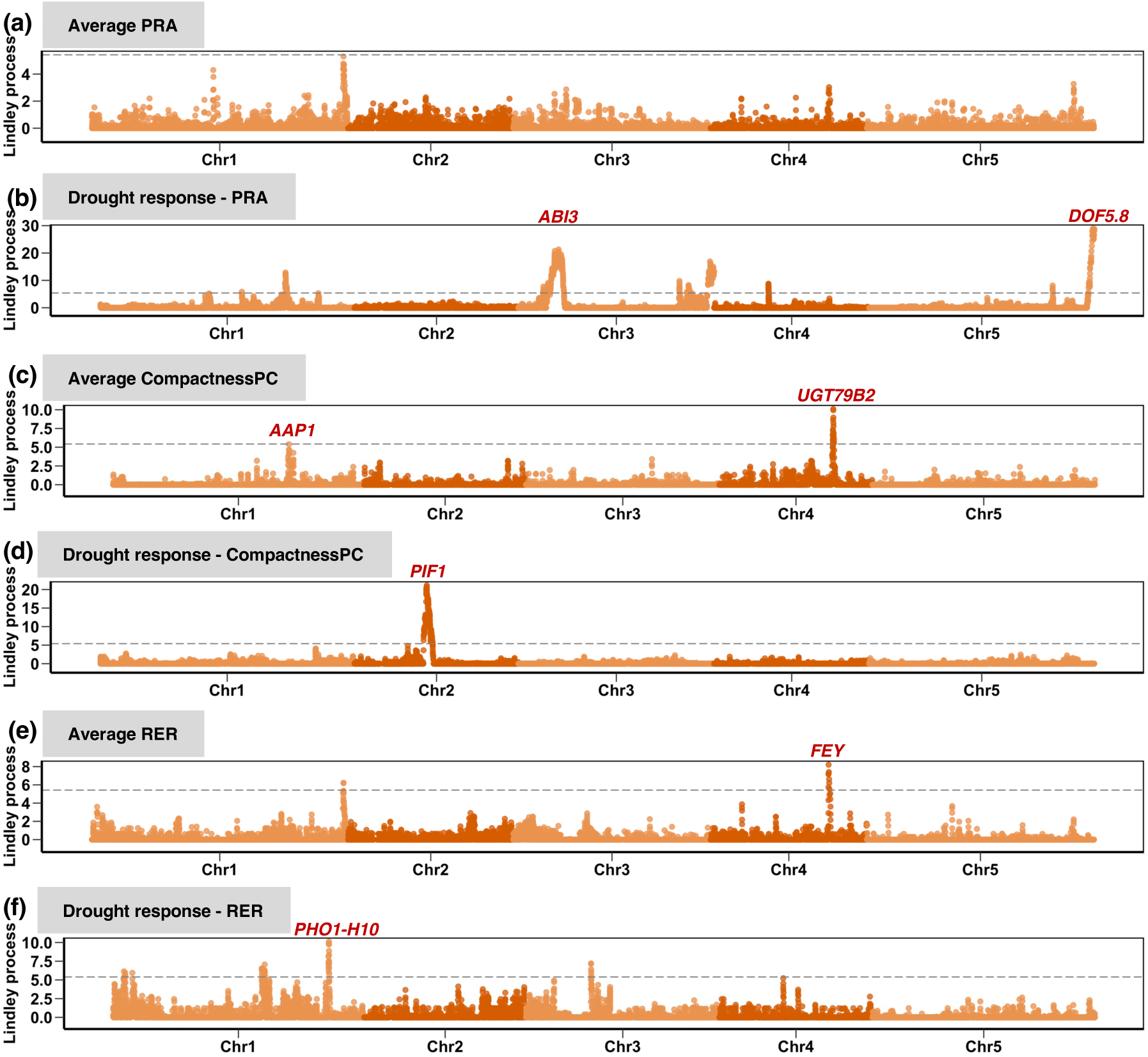
Genomic associations of rosette growth-related traits (PRA, compactnessPC, and RER) in Fogo Antão using LMM in GEMMA followed by the local score approach. The x-axis and y-axis represent the chromosomes and Lindley score from the local score approach. The dashed line in plots corresponds to a genome-wide Bonferroni significance.

**Figure S7.**
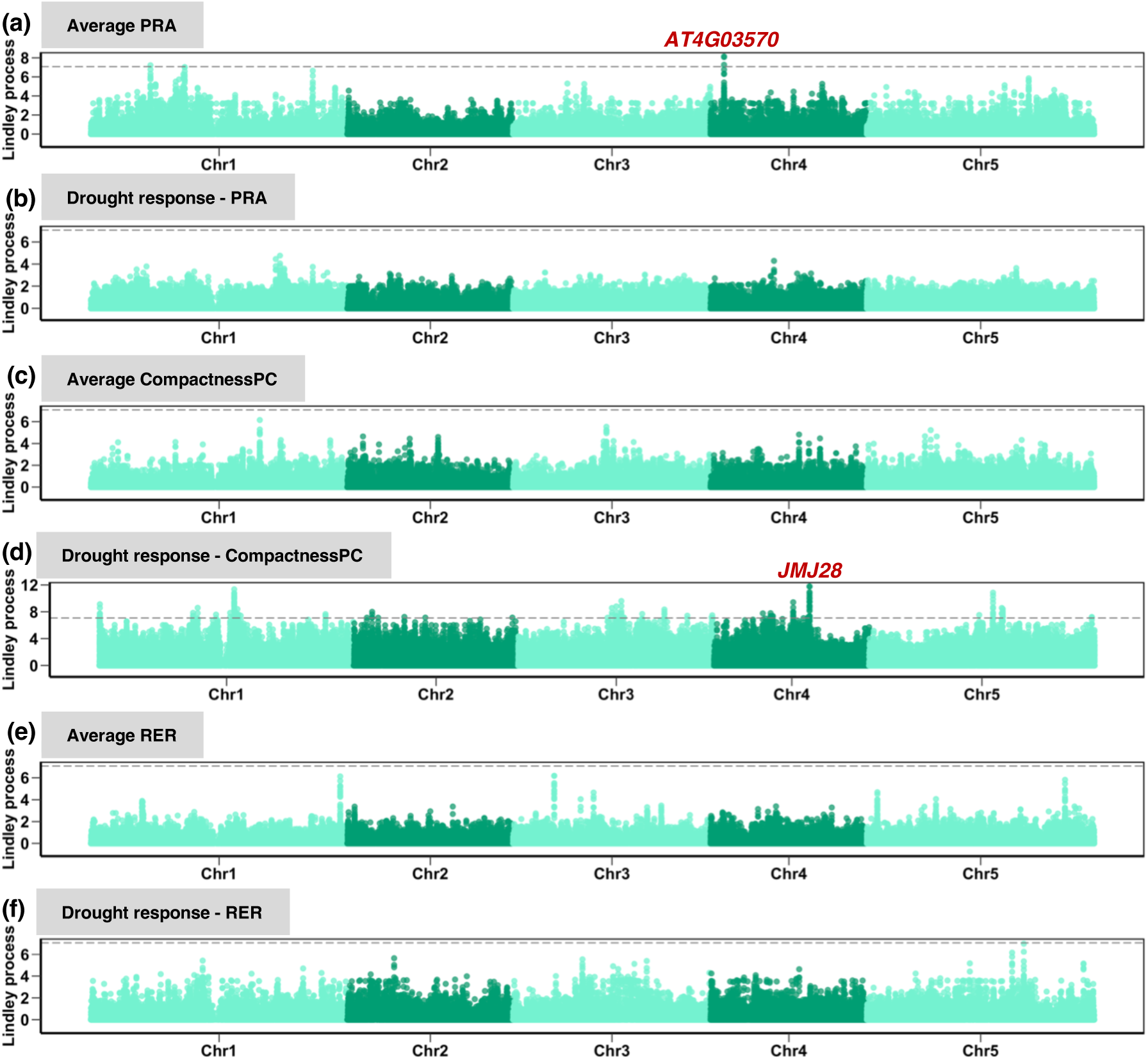
Genomic associations of rosette growth-related traits (PRA, compactnessPC, and RER) in Morocco using LMM in GEMMA followed by the local score approach. The x-axis and y-axis represent the chromosomes and Lindley score from the local score approach. The dashed line in plots corresponds to a genome-wide Bonferroni significance.

**Figure S8.**
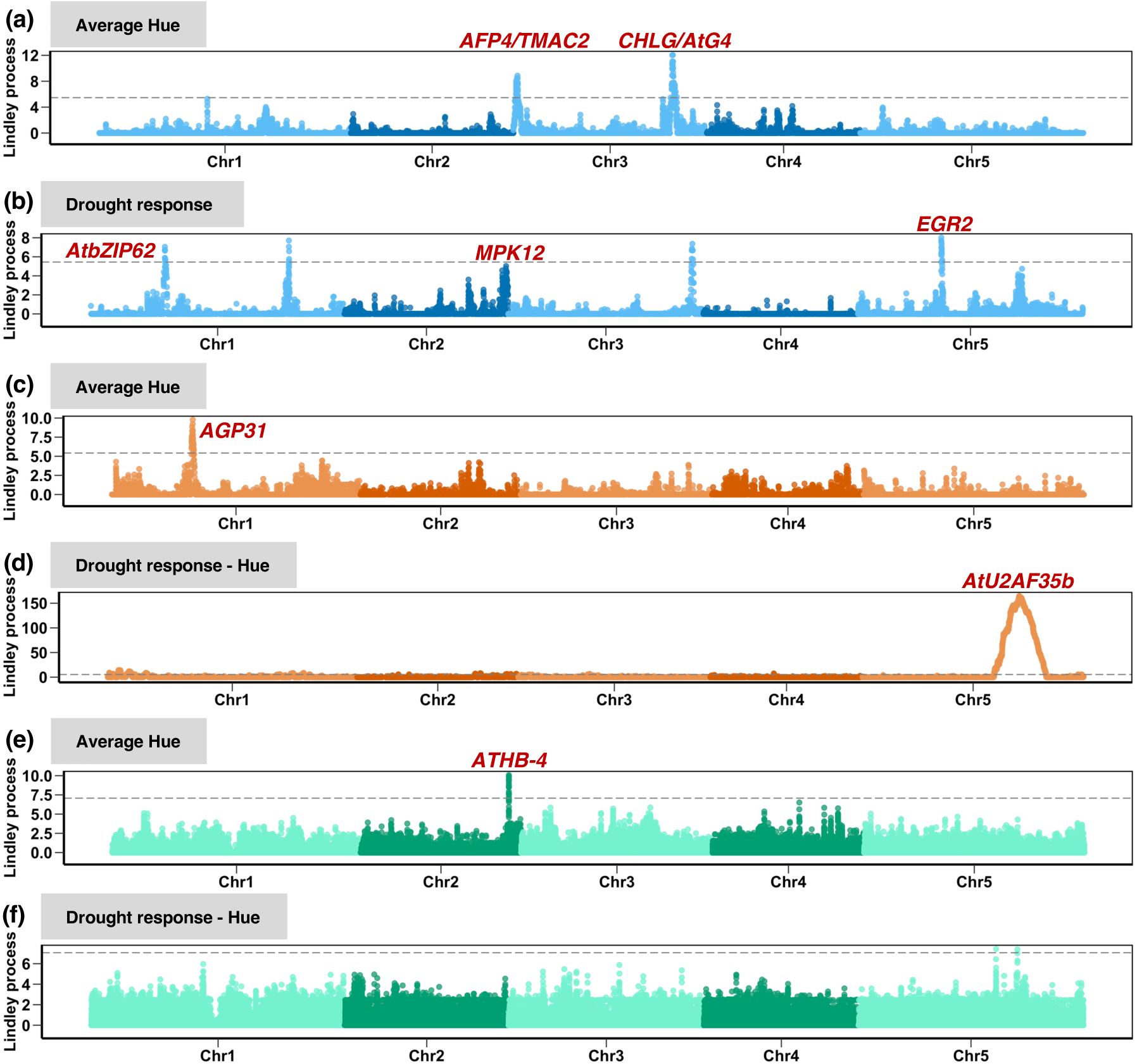
Genomic associations of hue (a proxy for rosette pigments content, including chlorophyll) in response to drought in Santo Antão (blue), Fogo (orange), and Morocco (green) populations using LMM in GEMMA followed by the local score approach. The y-axis represents the Lindley process score from the local score approach. The dashed line in plots corresponds to a genome-wide Bonferroni significance.

**Figure S9.**
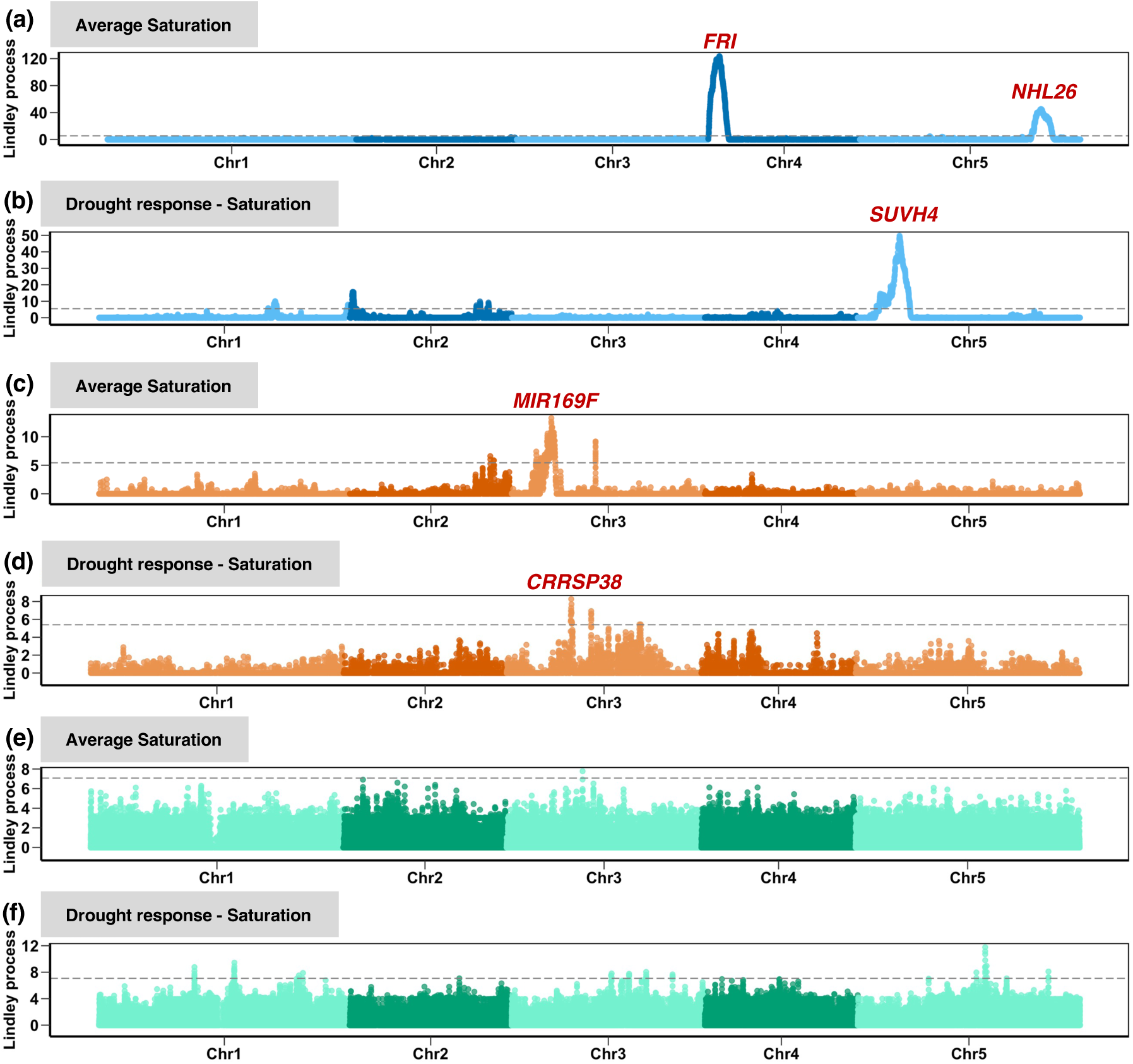
Genomic associations of saturation (rosette color intensity) in response to drought in Santo Antão (blue), Fogo (orange), and Morocco (green) populations using LMM in GEMMA followed by the local score approach. The y-axis represents the Lindley process score from the local score approach. The dashed line in plots corresponds to a genome-wide Bonferroni significance.

## Notes

### Competing Interest Statement

The authors have declared no competing interest.

